# Modelling the Decay of Hotspot Motifs in Broadly Neutralizing Antibody Lineages

**DOI:** 10.1101/055517

**Authors:** Kenneth B Hoehn, Gerton Lunter, Oliver G Pybus

**Affiliations:** Department of Zoology, University of Oxford, South Parks Road, Oxford, OX1 3PS, UK; Wellcome Trust Centre for Human Genetics, University of Oxford, Roosevelt Drive, Oxford, OX3 7BN, UK

## Abstract

Phylogenetic methods have shown great promise in understanding the development of broadly neutralizing antibody lineages (bNAbs). However, mutational process for generating these lineages - somatic hypermutation (SHM) - is biased by hotspot motifs, which violates important assumptions in most phylogenetic substitution models. Here, we develop a modified GY94-type substitution model which partially accounts for this context-dependency while preserving independence of sites in calculation. This model shows a substantially better fit to three well-characterized bNAb lineages than the standard GY94 model. We show through simulations that accounting for this can lead to reduced bias of other substitution parameters, and more accurate ancestral state reconstructions. We further explore other implications of this model; namely, that the number of hotspot motifs - and therefore likely the mutation rate in general - is expected to decay over time in individual bNAb lineages.

## Introduction

Recent advances in sequencing technology are giving an unprecedented view into the genetic diversity of the immune system during infection, especially for chronic infections caused by viruses. Broadly neutralizing antibody (bNAb) lineages, which produce B cell receptors (BCRs) capable of binding a wide range of viral epitopes, are of particular interest (Haynes et al. 2012). Within such lineages, all B cells descend from a shared common ancestor and are capable of rapid sequence evolution through the processes of somatic hypermutation (SHM) and clonal selection. For chronically infecting viruses such as HIV–1, this co-evolutionary process may continue for years (Wu et al. 2015). Because immunoglobulin gene sequences from bNAb lineages undergo rapid molecular evolution, selection and diversification, they would appear to be suitable for evolutionary and phylogenetic analysis, and these methods have already been applied to various immunological questions such as inferring the ancestral sequences of bNAb lineages (Sok et al. 2013; Hoehn et al. 2016). These intermediate ancestors are of particular interest because they may act as targets for stimulation by vaccines (Haynes et al. 2012).

However, the biology of mutation and selection during somatic hypermutation is different from that which occurs in the germline, and therefore it is unlikely that standard phylogenetic techniques will be directly applicable to studying bNAb lineages without suffering some bias and error. One of the most important assumptions of likelihood-based phylogenetics is that evolutionary changes at different nucleotide or codon sites are assumed to be statistically independent. Without this assumption, likelihood calculations rapidly become computationally impractical as the length and number of sequences increases (Felsenstein 1981). Unfortunately, in contrast to germline mutations, somatic hypermutation of BCR sequences is driven by a specific enzyme, called activation induced cytodine deaminase (AID), which targets defined “hotspot” sequence motifs that are usually 2- 3bp long (Yaari et al. 2013). This specificity clearly violates the assumption of independent evolution among sites. Furthermore, because hotspot motifs are, by definition, more mutable than non-hotspot motifs, we propose that their frequency within a B-cell lineage may decrease over time as they are replaced with more stable motifs. These changes will not be inherited because the mutational process is somatic. This effect may have a number of consequences on molecular evolutionary inference, for example it may render inappropriate the common practice of estimating equilibrium frequencies from the sequences themselves. At present it is unknown how the violation of these assumptions might affect phylogenetic inference of BCR sequences in practice, and the problem of ameliorating such effects remains an open issue.

This work has two main aims. The first is to analyse BCR evolution in three previously-published and long-lived bNAb lineages in HIV–1 infected patients. This analysis confirms our prediction of a decay of certain hotspot motifs through time. Our second aim is to develop and introduce a new substitution model that can partially account for this effect. The model is a modification of the GY94 (Goldman and Yang 1994) codon substitution model. Although only an approximation, our new model can detect and quantify the effect of AID-mediated somatic hypermutation on BCR sequences whist preserving the assumption of independence among codon sites to maintain computational feasibility. This model shows a significantly better fit than the standard GY94 model to all three bNAb lineages from HIV-1 patients. Through simulations, we further show that this model reduces bias in the estimation of other evolutionary parameters we and explore its potential applications, such as improved ancestral state reconstruction.

## Methods

### Multiple Sequence Alignment

Heavy chain sequences from the three bNAb lineages presented in (Wu et al. 2015) were downloaded from GenBank (http://www.ncbi.nlm.nih.gov/genbank/). The lineage of greatest duration was VRC01, which was sampled over 15 years (Wu et al. 2015), followed by CAP256, which was sampled over four years (Doria-Rose et al. 2014), and CH103, which was sampled over three years (Liao et al. 2013). Sequences from each bNAb lineage were translated into amino acids, aligned to their putative germline V gene segment using IgBlast (Ye et al. 2013), and then re-translated back into codons. Putative germline segment assignments (V4-59^*^01 for CH103, V3-30^*^18 for CAP256, and V1-2^*^01 for VRC01) were obtained from bNAber (Eroshkin et al. 2013) and sequences were obtained from the IMGT V-Quest human reference set (Lefranc and Lefranc 2001). Because of considerable uncertainty in D and J germline assignments for each lineage, only the V segment was used. Insertions relative to the germline sequence were removed, so that all sequences within each lineage were aligned to the same germline sequence. When this was done, the 3’ nucleotide of the region joined together from the removal of the insertion was converted into an N to avoid creating false motifs. To keep results consistent between lineages, only nucleotide positions from the beginning of the first framework region (FWR1) to the end of FWR3 were used. Sampling dates of each sequence were extracted from the sequence ID tags provided on GenBank. When these were not available the sequence was excluded.

### Hotspot decay in bNAb lineages

The “hotspot frequency” of each sequence was defined as the number of hotspot motifs divided by the possible number of hotspot motifs in that sequence, and was calculated for trimer (WR<u>C</u>/<u>G</u>YW) and dimer (W<u>A</u>/<u>T</u>W) motifs separately (Yaari et al. 2013), where W = A or T, Y = A or G, and R = T or C, as per the IUPAC nucleotide ambiguity codes. Hence an example trimer motif might be AT<u>C</u>, and its reverse complement <u>G</u>AT. The underlined base in each of these motifs experiences increased AID mediated mutability. Trimers and dimers with non-ACGT characters were excluded from this calculation. Changes in hotspot frequency values through time were analysed using linear regression and correlation. Because the date of infection was not known for VRC01, germline IGHV sequences were not included in these calculations. Importantly, because the sequences within each B-cell lineage are phylogenetically related, they are partially correlated due to shared common ancestry and are not independent data points, hence p-values from standard correlation and regressions tests are not reliable. However, the regression can still be an unbiased measure of trends in sequence change over time (see (Drummond et al. 2003) for discussion). Regressions of hotspot frequency through time are shown in Figure 1.

**Figure 1:**
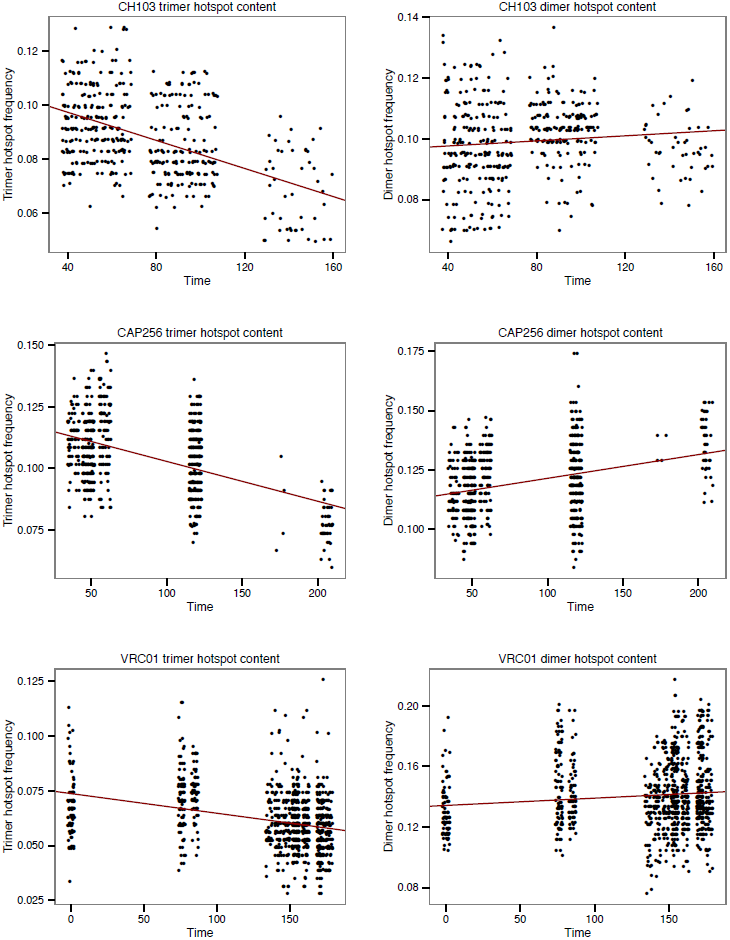
Decrease in frequency of trimer and dimer hotspot motifs in three bNAb lineages. Red line shown is least square regression.

In the absence of a suitable hypothesis test based on regression, we developed a simulation-based significance test for the association between hotspot frequency and time in bNAb lineages. The null model for this test is a substitution model (GY94) that does not explicitly model the decay of hotspot motifs and which is used to estimate a maximum likelihood phylogenetic tree. Multiple data sets were simulated under this null model, using the same sample sizes and sampling times as the three empirical bNAb data sets. The significance of the difference between the null model and the observed data is calculated as the proportion of simulated datasets with a
greater negative correlation between hotspot frequency and time than in the observed data set. Results for these tests are shown in Table 1.

**Table 1:** Hotspot motif decay in three bNAb lineages.

| B-cell lineage | Trimer motifs: WRC/GYW |  |  | Dimer motifs: WA/TW |  |  |
| --- | --- | --- | --- | --- | --- | --- |
|  | Observed correlation | Observed/simulated | P value | Observed correlation | Observed/simulated | P value |
| CH103 | -0.48 | -11.33 | 0.00 | 0.09 | 0.46 | 0.29 |
| CAP256 | -0.50 | -11.77 | 0.00 | 0.33 | 0.84 | 0.30 |
| VRC01 | -0.33 | 5.50 | 0.02 | 0.11 | 0.70 | 0.39 |

The “Observed correlation” column shows the correlation between hotspot frequency and time. The next column shows how these values compare to the mean of the same values from 100 simulations under the null model. The third column shows the p value – the proportion of simulated data sets that had a lower correlation than observed data sets.

Maximum likelihood phylogenetic trees and substitution model parameters for each of the three bNAb lineages were estimated using the GY94 model and empirical codon frequencies using codonPhyML (Gil et al. 2013). Trees were re-rooted so as to place the germline sequence as an outgroup with a branch length of zero, effectively making it the universal common ancestor (UCA) of the lineage. For each bNAb lineage, we then simulated 100 sequence data sets down the corresponding ML tree using the GY94 model, starting with the corresponding germline sequence at the root and using the fitted substitution model parameters. Simulations were performed using the PAML evolver program (Yang 2007).

To ascertain whether the observed effects were general, or specific to known hotspot motifs, we repeated the above regression and simulation tests for non-hotspot motifs. To do this, we randomly assigned non-hotspot nucleotide motifs as “hotspots” whilst keeping the number of trimer and dimer hotspots the same (eight and three, respectively). This analysis was then repeated for 100 such random allocations. These results are summarized in **Supplemental File 1**.

### A Codon Substitution Model for Phylogenies Undergoing Somatic Hypermutation

In order to represent the molecular evolution of long-lived B cell lineages more accurately, we develop a new substitution model that models the effects of motif-specific mutation across the BCR. This model, named the HLP16 model, is a modification of the GY94 substitution model. Specifically, we add to the GY94 model an additional parameter, *h*, which represents the additional increase in relative substitution rate of a hotspot mutation. Explicitly modelling the full context dependence of hotspot motifs would make likelihood calculations computationally infeasible. Instead, we weight *h* by *b_ij_*, which is the probability that the mutation from codon *i* to codon *j* was a hotspot mutation, averaged across all possible combinations of codons on the 5’ and 3’ flanks of the target codon. A “hotspot mutation” is defined as a mutation occurring within the underlined base of the trimer motif WR<u>C</u>/<u>G</u>YW. Because we did not find a significant decay of dimer hotspot motifs through time (**see** Figure 1 **and** Table 1), our model only includes trimer hotspots. However, dimers or other motifs could easily be added with additional values of *h* and *by* for each new motif.

In the HLP16 model, each entry q_ij_ in the Markov rate matrix Q is defined by:

π_j_: Equilibrium frequency of codon j
k: Transition/transversion mutation relative rate ratio
ω: nonsynonymous to synonymous mutation relative rate ratio
h: Increased mutability due to trimer hotspot motifs (WR<u>C</u>/<u>G</u>YW)
b_ij_: Probability that mutation from *i* to *j* occurred in a trimer hotspot

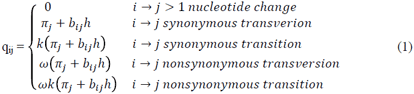

The values of *b*_ij_ are calculated by marginalizing over all possible 5’ and 3’ flanking sense codons as follows:

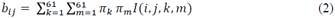

Where *I* is the indicator function:

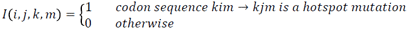

This model, though an approximation, has several useful properties. Perhaps most importantly, because codon changes are still modelled independently of each other, likelihood can still be calculated using Felsenstein’s pruning algorithm, which greatly improves computational time (Felsenstein 1981). The model also has the intuitive property that, if hospots are subjected to no additional mutability compared to other motifs, then *h* = 0 and the model simplifies to the GY94 model. Thus the GY94 model is a special case of the HLP16 model. Further, mutations that could have not occurred instantaneously within a hotspot motif reduce to their GY94 values before Q matrix normalization.

In contrast to most substitution models, the relative substitution rate parameters in the Q matrix of the HLP16 model are not necessarily symmetric. A Q matrix with symmetric rate parameters is useful because it likelihood calculations can be undertaken on an unrooted tree, which can then be rooted by any of its tips. In the case of B cell lineage evolution, it is necessary to root the lineage phylogeny at the germline sequence during parameter optimization, and pruning algorithm likelihood calculations are required to always begin at the root node.

We modified the source code of codonPhyML (Gil et al. 2013) to implement the rate matrix in equation 1, and to estimate the additional parameter *h* using maximum likelihood, alongside the other model parameters. Specifically, we optimize ω, k, π_j_ (using the CF3X4 model) and the vector of phylogeny branch lengths. Performing all likelihood calculations from the root node slowed computation substantially, therefore in this work we applied the HLP16 model to a fixed tree topology, and we leave the problem of co-estimating topology for future work. For each data set, the tree topology used was that inferred using the standard GY94 model in codonPhyML, which was subsequently re-rooted to place the germline sequence at the universal common ancestor.

Because the GY94 model is a special case of the HLP16 model (i.e. when *h* = 0) the two models are nested and can be compared using a likelihood ratio test. Let *max L(HLP16)* and *max L(GY94)* be the maximum likelihoods obtained under the HLP16 and GY94 models, respectively. The likelihood ratio statistic is then 2^*^( *max L(HLP16)* / *max L(GY94)* ) and approximately chi-squared distributed with one degree of freedom (Huelsenbeck and Rannala 1997). For each bNAb dataset, the value *max L(HLP16)* is calculated by optimising *h* together with the other model parameters, whereas the value *max L(GY94)* is calculated by constraining *h*=0 whilst optimising all other model parameters.

We evaluated the performance of the HLP16 model by simulating data sets under different values of *h* and testing how accurately *h* and other model parameters were inferred. The following procedure we used to generate the simulated data sets:

1. We randomly subsampled each bNAb lineage to 99 sequences, plus the single germline sequence at the root. Subsampling was necessary to make the large number of replicates computationally feasible.
2. We estimated a maximum likelihood phylogeny for each subsampled bNAb lineage data set using the standard GY94 model. During estimation we optimised ω, k, π_j_, branch lengths and the tree topology. The resulting ML tree was re-rooted at the germline sequence with a branch length of zero.
3. For each value of *h* investigated (0, 0.05, 0.1, and 0.3), we simulated 10 alignments along each of these trees. Simulations were undertaken using the estimated values of ω, k and π_j_, obtained in step (2) for the corresponding bNAb lineage datra set. Starting (root) sequences were generated randomly from equilibrium frequencies.
4. For each of the replicates defined in step (3), we performed three different ML calculations: (i) *h* was optimised using ML (with *ĥ* as the MLE estimate of *h*), (ii) *h* was fixed to zero, and (iii) *h* was fixed to the true value used in simulation *(h_true_).* These three values enable us to test both type 1 and type 2 error rates, by determining whether *ĥ* was significantly different to *h_true_* or *h_0_* respectively. Statistical significance was determined using the chi-squared approximation to the likelihood ratio statistic, as described above. In all calculations, the tree topology was fixed to that inferred from step (2).
5. For each data set and for each set of simulations under a particular value of h, we estimated *h* and then calculated the properties of this estimator as follows:

i. Bias in estimation: (Mean[*ĥ*] – *h_true_*)
ii. Variance in estimation: Variance[*ĥ*]
iii. Type 1 error rate: The proportion of simulated data sets in which *h_true_* was outside of the 95% confidence interval for *ĥ*.
iv. Type 2 error rate: The proportion of simulated data sets in which *h >*0, but failed to reject the null hypothesis (*h_0_*).

Biased mutation during somatic hypermutation has been shown to give false signatures of natural selection using approaches that compare the expected number of replacement and silent mutations under a null model of somatic hypermutation (Dunn-Walters and Spencer 1998), and it is possible that the HLP16 model could partially reduce this bias. To test this, and to explore whether the HLP16 model improved the estimation of other evolutionary parameters, we compared the percentage error under the HLP16 and GY94 models of estimates of (i) ro, (ii) k, (iii) tree length (sum of all branch lengths) and (iv) the ratio of internal to external branch lengths. These results are provided in **Supplemental Figure 2.**

The fact that bNAb lineages are clearly not in equilibrium when they are sampled (Figure 1) has interesting implications for the use of Markov substitution models. Typically, it is assumed that nucleotide or codon frequencies are at equilibrium at the time of sampling, and empirical codon frequencies are often to estimate equilibrium frequencies. In the case of long-lived B cell lineages, however, sampled sequences are almost certainly not in equilibrium and do not converge to an equilibrium because the changes are somatic and not inherited, thus making empirical codon frequencies inaccurate. For this reason, it is necessary to optimize equilibrium frequency using ML rather than use empirical codon frequencies. To test how this might affect estimation of *h*, we repeated the simulation procedure above using empirical equilibrium frequencies from each data set. These results are included in **Supplemental File 3**.

Because the HLP16 model is a mean field approximation to context dependency of hotspot mutations, it is unlikely to fully account for the context dependency of somatic hypermutation. To test how this may affect analyses, we repeated our simulation procedure using a model that fully accounts for the context dependence of adjacent codon sites. In this forward simulation procedure, because the 3’ and 5’ flanking codons of each site are known, we create a B matrix for each site in each sequence with *b_ij_* equal to either 1 or 0 depending on whether the substitution was or was not in a hotspot mutation. This process begins at the root sequence, calculates a separate Q matrix at each site in the sequence, and simulates two descendant sequences down the tree until all tips are filled.

One of the key application of molecular phylogenetics to BCR sequence data is the reconstruction of ancestral sequences within a B-cell lineage (Kepler 2013). Ancestral state reconstruction is an implicit part of the phylogenetic likelihood calculation when nucleotide or codon substitution models are used. For each simulation replicate, and for each of the three likelihood calculations described in step (3) above, we computed the most likely codon at each codon position at each internal node in the tree. These ancestral sequences were then used to compare the accuracy of reconstructions under the HLP16 model with those obtained using the GY94 model. In each simulation replicate, accuracy of ancestral sequence reconstruction was measured by calculating the mean pairwise nucleotide or amino acid difference between the predicted and true sequences at each node.

## Results

### Decay of hotspot motifs in bNAb lineages

All three bNAb lineages showed a negative correlation between trimer hotspot content and time. However, no such decline was seen in dimer motifs (Table 1, Figure 1). To test whether the patters of hotspot decay observed were significantly different from those expected under a standard reversible codon substitution model that does not explicitly account for hypermutation at hotspot motifs, we implemented a significance test that compares the correlation between hotspot motif frequency and time in simulated data sets generated under the null phylogenetic model. All three B cell lineages showed a significantly greater negative correlation between trimer hotspot content and time than expected under the null model (Table 1). In all cases, the frequency of dimer motifs showed no significant change through time. Furthermore, we repeated these analyses with randomly-chosen non-hotspot motifs taking the place of the real, known hotspot motifs. This latter analysis demonstrates that the significant decline detected was specific to known hotspot motifs, as declines of similar degree were rarely observed in non-hotspots (**Supplemental File 1**).

### A codon substitution model for phylogenies undergoing somatic hypermutation

All three bNAb lineages showed a significant improvement in likelihood under the HLP16 model compared to the GY94 model. The maximum likelihood values of *h* for the three data sets were *ĥ* = 0.0345, 0.032, and 0.0339, for CH103, CAP256, and VRC01, respectively. In each case the simpler GY94 model *(h=0)* could be rejected using the likelihood ratio test (*p* < 1×10^−5^ for all three lineages). These results are summarized in Table 2. While the absolute values of *ĥ* may appear small, it is important to remember that *h* is added to the equilibrium frequencies before being multiplied by other factors (see equation 1). Because the average equilibrium frequency is approximately 1/61 = 0.016, a *h* value of 0.034 represents up to a threefold increase in the relative rate for hotspot mutations (depending on the values of *b_ij_*).

**Table 2:** Maximum likelihood estimates of *h* and likelihood ratio tests

| B-cell lineage | $Mean \hat{h}$ | Log-likelihood | | $2*LR$ | p value |
| --- | --- | --- | --- | --- | --- |
| | | $h_{MLE}$ | $h_0$ | | |
| CH103 | 0.0345 | -14918.9 | -15031.5 | 225.2 | $<1 \times 10^{-50}$ |
| CAP256 | 0.0322 | -37622.1 | -37913.8 | 583.4 | $<1 \times 10^{-50}$ |
| VRC01 | 0.0339 | -44011.2 | -44339.3 | 656.2 | $<1 \times 10^{-50}$ |

Significance was determined using the likelihood ratio test under a chi-squared distribution with one degree of freedom.

Our implementation of the HLP16 mode proved to be a robust ML estimator of *h* when it was applied to simulated bNAb lineages (Table 3). Mean *ĥ* values were very close to their true *h* values, with low bias (maximum 9.3×10^−3^) and variability (maximum 7.6×10^−4^). None of the 120 data sets simulated with *h*? > 0 failed to reject the null hypothesis that *h* = 0. Further, our method incorrectly rejected the true value of *h* in 4.2% of analyses, which is approximately as expected given an alpha value of 0.05. Performance of inference was highly consistent across the three bNAb phylogenies used to generate simulated alignments. We found that using empirical equilibrium frequencies decreased the efficiency of parameter estimation, with higher bias and type 2 error rates than when ML equilibrium frequencies were used. See Methods for some discussion of why empirical codon frequencies are unlikely to be suitable for long-lived B-cell lineage phylogenies (**Supplemental File 3**).

**Table 3:**
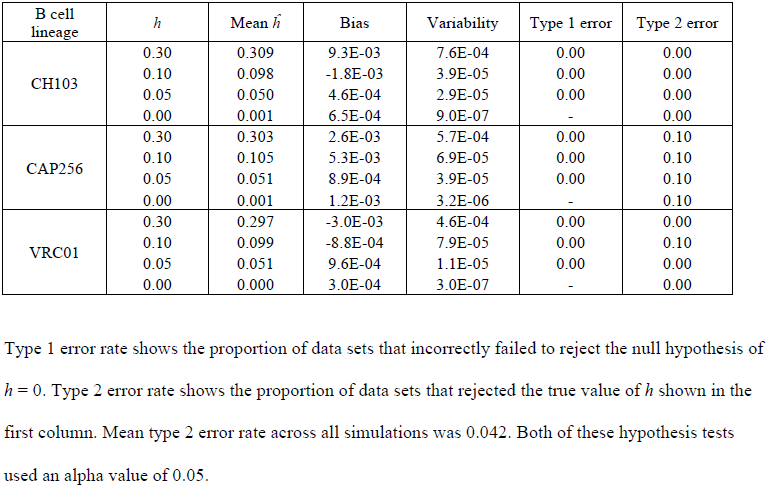
Analyses of data sets simulated using different phylogenies and values of *h*

| B cell lineage | $h$ | Mean $\hat{h}$ | Bias | Variability | Type 1 error | Type 2 error |
| --- | --- | --- | --- | --- | --- | --- |
| CH103 | 0.30 | 0.309 | 9.3E-03 | 7.6E-04 | 0.00 | 0.00 |
|  | 0.10 | 0.098 | -1.8E-03 | 3.9E-05 | 0.00 | 0.00 |
|  | 0.05 | 0.050 | 4.6E-04 | 2.9E-05 | 0.00 | 0.00 |
|  | 0.00 | 0.001 | 6.5E-04 | 9.0E-07 | - | 0.00 |
| CAP256 | 0.30 | 0.303 | 2.6E-03 | 5.7E-04 | 0.00 | 0.10 |
|  | 0.10 | 0.105 | 5.3E-03 | 6.9E-05 | 0.00 | 0.10 |
|  | 0.05 | 0.051 | 8.9E-04 | 3.9E-05 | 0.00 | 0.10 |
|  | 0.00 | 0.001 | 1.2E-03 | 3.2E-06 | - | 0.10 |
| VRC01 | 0.30 | 0.297 | -3.0E-03 | 4.6E-04 | 0.00 | 0.00 |
|  | 0.10 | 0.099 | -8.8E-04 | 7.9E-05 | 0.00 | 0.10 |
|  | 0.05 | 0.051 | 9.6E-04 | 1.1E-05 | 0.00 | 0.00 |
|  | 0.00 | 0.000 | 3.0E-04 | 3.0E-07 | - | 0.00 |

Type 1 error rate shows the proportion of data sets that incorrectly failed to reject the null hypothesis of $h = 0$ . Type 2 error rate shows the proportion of data sets that rejected the true value of $h$ shown in the first column. Mean type 2 error rate across all simulations was 0.042. Both of these hypothesis tests used an alpha value of 0.05.

Incorporating and estimating the parameter *h* in the HLP16 model appears to have generally improved the estimation of other model parameters, in comparison to the GY94 model (Figure 2, **Supplemental File 2**). The clearest improvement was seen in the tree length parameter; when *h* is large, the GY94 model substantially underestimates tree length for simulations based on the VRC01 lineage phylogeny (mean percentage error up to -0.4) and there are clear biases for the simulations based on other phylogenies (**Supplemental File 2**) as well. The bias was effectively absent when *h* = 0 and increased as *h* increased. In contrast, the HLP16 model did not lead to underestimation of tree length as *h* increases. Analyses of simulated data sets also showed that in some cases the GY94 model progressively overestimates the ro parameter as the true value of *h* increases; this bias was clear in simulations based on the most diverse B-cell lineage (VRC01) but was not obvious in simulations based on the CAP256 and CH103 lineages that were sampled for a shorter duration. Again, the HLP16 model was able to estimate ro accurately under all values of *h*.

**Figure 2:**
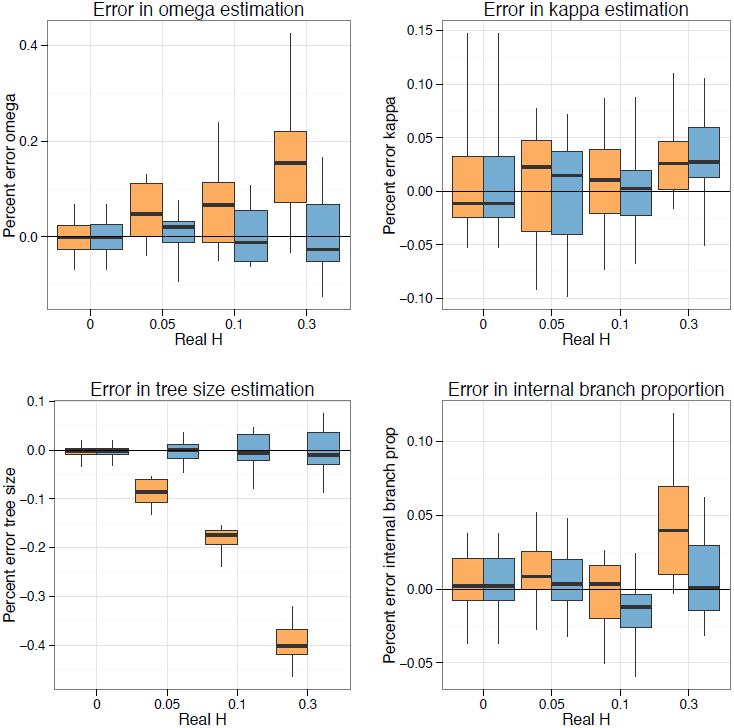
Percentage error in parameter estimation compared to true values for the VRC01 B cell lineage. Estimates obtained under the GY94 are in orange (*h*=0) and estimates obtained under the HLP16 model are in blue (*h* estimated using maximum likelihood). The edges and centers of boxplots show the 1^st^, 2^nd^, and 3^rd^ quartiles, while the whiskers show range. Equivalent results for B cell lineages CH103 and CAP256 are shown in **Supplemental File 2**.

Repeating our simulation analysis on data sets simulated under a model that fully accounts for the context dependence of hotspot mutations showed expected decreases in accuracy relative to simulations under the approximate model. Inference of *ĥ* was generally underestimated in simulations where *h* > 0, and statistics such as tree size and internal/external branch length ratios were biased as well under large values of h. However, the HLP16 model showed consistently equal or better performance in all of these categories compared to the standard GY94 model, showing both a significantly higher likelihood in all data sets in which *h* >? 0, and generally equal or better inference of other parameters and tree statistics. Importantly, these biases and losses in accuracy were primarily seen for large values of *h* (> 0.05), many times larger than the *ĥ* values we inferred from bNAb lineages. Further, values of *h* tended to be underestimated in these simulations, indicating that our estimates of *ĥ* from bNAb lineages are likely conservative.

We also found that the HLP16 model provided more accurate reconstructions of ancestral sequences than the GY94 model in most simulations (**Supplemental File 4**). As expected, the level of improvement increased as the true value of *h* increased.

Sequence reconstructions under the two models were fairly similar for internal nodes near the root and the tree tips, and the improvement under the HLP16 model was most marked for internal nodes in the basal third of the phylogeny. Typically, we would expect the uncertainty in ancestral state reconstruction to increase as we move from the tree tips towards the root, but B-cell lineages are unusual in that the root sequence is also known as it corresponds to the germline sequence.

While true ancestral sequences are not available for the three bNAb lineages, ancestral state reconstructions we did observe differences between sequences reconstructed from HLP16 and GY94 models. For each lineage we compared these models by calculating the mean amino acid difference between the predicted ancestral sequences at all internal nodes of each tree. Performing this ancestral state reconstruction on each of the three bNAb lineages showed a mean of 0.31, 0.62, and 0.35 amino acid sequence difference across all internal nodes with most differences also concentrated near the upper-middle of the tree (**Supplemental File 4**).

## Discussion

Molecular phylogenetics has already been used in a variety of applications in the study BCR genetic diversity and the molecular evolution of B cell lineages (Kepler 2013; Sok et al. 2013; Kepler et al. 2014). However, the process of somatic hypermutation is known to operate in ways that violate fundamental assumptions of most phylogenetic substitution models. Here, we demonstrate that failing to account for the violation of these assumptions has tangible effects on phylogenetic inference and parameter estimation from sets of sequences from long-lived bNAb lineages. We develop and implement a new codon substitution model (HLP16) that, whilst only an approximation, is capable of mitigating these effects.

Perhaps the most salient difference between standard substitutions models and the biology of somatic hypermutation is the context dependency of mutation in BCRs. This has long been known to give false signature of selection in BCRs (Dunn-Walters and Spencer 1998). This effect was observed in our own simulations (**Supplemental File 2**) in which failing to account for the increased rate of substitution at hotspot motifs led to overestimation of the ro (dn/ds) parameter. A variety of empirical models have been developed to characterize biased hotspot motifs at di, tri, penta, and heptamer levels (Smith et al. 1996; Yaari et al. 2013; Elhanati et al. 2015).

Some approaches have been developed to study the substitution process in BCR data in the context of biased mutation. Some of these are non-phylogenetic in nature (e.g. Hershberg et al. 2008; Yaari et al. 2012) and focus on the expected number of germline to tip replacement mutations regions in comparison to a null model. Kepler et al (Kepler et al. 2014) developed a non-linear regression model approach that, combined with an empirical model of mutation rate at each site, allowed the authors to test for both selection, and the interaction between mutation and selection, in shaping BCR genetic diversity. The substitution model detailed in McCoy et al (McCoy et al. 2015) is perhaps the most similar to the new model introduced in this study, as it is ultimately derived from probabilistic reversible substitution models. This was accomplished by comparing values of ro inferred from a given data set to those inferred from out of frame rearrangements.

The HLP16 codon substitution model detailed here is a relatively straightforward modification of the widely used GY94 model. Although the HLP16 model is slower to compute than the simpler, reversible model on which it is based, we have found that it is usable, and almost certainly statistically preferable, to the GY94 model when applied to any BCR data set whose diversity may have been shaped by somatic hypermutation. Further, the HLP16 model does not rely on an empirical model to incorporate the effect of biased mutation, but instead attempts to explicitly model the context-dependent mutational process by estimating the parameter *h* and equilibrium codon frequencies directly from the data. It should be noted, however, that the HLP16 model is a mean-field approximation and does not capture the full context of motif driven evolution. Therefore we do not expect it to fully disentangle interaction between selection and biased mutation, and estimated values of ro should be interpreted carefully. In addition to correcting biases in parameter estimation, simulation analyses reveal that the HLP16 model produces different, and more accurate, ancestral state reconstructions than the standard GY94 model.

Another common assumption in phylogenetic analysis is that the codons or nucleotides sampled for analysis are at their equilibrium frequencies. Because our hotspot model has asymmetric relative rates between codons, which are a function of h, then it seems likely that codon frequencies will change across a B-cell lineage when *h* is significantly above zero. This is a result of the decline in the number of hotspots through time (Figure 1). We dealt with this problem by estimating equilibrium frequencies by maximum likelihood within the model. This showed improvement, both in maximum likelihood values and in parameter estimation, over using empirical codon frequencies to approximate equilibrium frequencies. However, it is not yet clear if this is the most efficient or the most effective way of dealing with this problem of sequence disequilibrium.

We suggest that this decay of hotspot motifs in bNAb lineages may have important implications for our understanding of host-virus coevolution. More specifically, we propose that the loss of hotspot motifs will lead to a decrease in sequence mutability, and therefore a decline in overall rate of evolution over time for a given lineage. Two important pieces of this hypothesis - lower mutability from loss of hotspot motifs, and decline in mutation rate over time in bNAb lineages - have already been explored separately. Wei et al. (2015) showed experimentally that a decrease in the number of hotspot motifs in a BCR sequence leads to a decrease in the overall mutation rate in that sequence. Further, a landmark study (Wu et al. 2015) showed multiple pieces of evidence that evolutionary rate in the same three bNAb lineages analysed here declines over time, a clear violation of standard clock rate models. The decay of hotspot motifs may at least partially explain this inferred slowdown in mutation rate. Whilst we do not directly test here the hypothesis that a decay of hotspot motifs over btime leads to decrease in overall BCR evolutionary rate, we do show that hotspot motifs decay within three bNAb lineages in HIV–1 infected individuals, which provides an empirical link between the ideas of biased mutation and the inferred decrease in substitution rate.

This hypothesis has an important corollary. If the slowdown in mutation rate over time, arising from hotspot decay, is an intrinsic property of activated B cell lineages, then BCR sequence divergence from a germline ancestor (and thus affinity maturation) must be fundamentally and intrinsically constrained. Consequently, while BCR lineages may be able to rapidly evolve binding affinity and co-evolve with pathogens for an initial period after activation, over longer periods of time B-cell lineages will fail to keep up with the rapid evolution of chronically infecting viruses such as HIV–1, due to the exhaustion of available BCR hotspot motifs. This would suggest using a single bNAb lineage against chronic viruses may be a losing strategy, and that utilizing multiple bNAb lineages (e.g. Gao et al. 2014) would be better suited to long-term coevolution with chronic viruses. From an evolutionary perspective, this hypermutation-driven “senescence” of B-cell lineages may be ultimately adaptive, because the fitness benefits of rapid short-term BCR sequence evolution accruing from acute infections would likely outweigh the costs of BCR hotspot motif exhaustion in the context of chronic infections.

Some of our findings are circumstantial support other hypotheses that may be interesting to test in future research. We observed that the relationship between hotspot frequency and time is stronger in the younger lineages (correlation = −0.48 and −0.5; for CH103 and CAP256), than in VRC01 (−0.33). This is consistent with the notion that the decline in the divergence rate of bNAb lineages is fastest early in the development of the B cell lineage, and slows down as the lineage ages. Interestingly, we returned a very similar estimates of *ĥ* (~0.03) for all three bNAb lineages tested, despite the fact that they had been evolved for different periods of time and other evolutionary parameter estimates (e.g. omega and kappa) varied among lineages. This may indicate that mutational bias arising from hotspot motifs is consistent across time scales and individuals.

Because our model conforms to certain limitations, such as independent change among codon sites, it cannot fully account for the effects of targeted SHM. Other properties of SHM are excluded in the model’s current form but may be possible to integrate. The most obvious property, and perhaps the easiest to accommodate, is the fact that other hotspots (e.g RGYW/WRCY, DGYW/ WRCH) and even cold spots (e.g. SYC) besides those explored here have been identified (Zhang et al. 2001; Rogozin and Diaz 2004; Peled et al. 2008). These may be accommodated by including a separate *h* parameter for each hotspot, and by modifying the indicator function to account for those hotspot or coldspot motifs. It is not clear how to best model the interaction between overlapping hotspot motifs, but both the inclusion of additional motifs, and the model of interaction between them may be tested through a likelihood ratio test. Another important issue is that BCR sequences are highly partitioned into framework (FWR) and complementary-determining (CDR) regions. These are known to be under different types and degrees of selection (Yaari et al. 2012; Yaari et al. 2015), so an obvious next step is to use a site-partition model to allow omega to vary between these two regions. While other analyses have already shown an interaction between region, mutability, and selection (Kepler et al. 2014), it
would be interesting to test this hypothesis in this framework by allowing *h* to vary between region partitions as well.

## Authors’ Note

After this manuscript was written, a paper by Sheng et al (2016) was published, on May 18^th^, which also explores the hypothesis that decline in mutation rate in bNAb lineages may be the result of hotspot motif loss. We were unaware of Sheng et al.’s manuscript whilst our work was undertaken.

Sheng Z, Schramm CA, Connors M, Morris L, Mascola JR, Kwong PD, Shapiro L. 2016. Effects of Darwinian Selection and Mutability on Rate of Broadly Neutralizing Antibody Evolution during HIV-1 Infection. PLOS Comput Biol 12:e1004940.

## Acknowledgements

This work was funded by the European Research Council under the European Union’s Seventh Framework Programme (FP7/2007-2013)/ERC grant agreement no. 614725-PATHPHYLODYN (to O.G.P.). KBH was funded through a Marshall scholarship.

## References

Doria-Rose NA, Schramm CA, Gorman J, Moore PL, Bhiman JN, DeKosky BJ, Ernandes MJ, Georgiev IS, Kim HJ, Pancera M, et al. 2014. Developmental pathway for potent V1V2-directed HIV-neutralizing antibodies. Nature 509:55–62.

Drummond A, Oliver G, Rambaut A, others. 2003. Inference of viral evolutionary rates from molecular sequences. Adv. Parasitol. 54:331–358.

Dunn-Walters DK, Spencer J. 1998. Strong intrinsic biases towards mutation and conservation of bases in human IgVH genes during somatic hypermutation prevent statistical analysis of antigen selection. Immunology 95:339.

Elhanati Y, Sethna Z, Marcou Q, Callan CG, Mora T, Walczak AM. 2015. Inferring processes underlying B-cell repertoire diversity. Phil Trans R Soc B 370:20140243.

Eroshkin AM, LeBlanc A, Weekes D, Post K, Li Z, Rajput A, Butera ST, Burton DR, Godzik A. 2013. bNAber: database of broadly neutralizing HIV antibodies. Nucleic Acids Res.:gkt1083.

Felsenstein J. 1981. Evolutionary trees from DNA sequences: A maximum likelihood approach. J. Mol. Evol. 17:368–376.

Gao F, Bonsignori M, Liao H-X, Kumar A, Xia S-M, Lu X, Cai F, Hwang K-K, Song H, Zhou T, et al. 2014. Cooperation of B Cell Lineages in Induction of HIV-1-Broadly Neutralizing Antibodies. Cell 158:481–491.

Gil M, Zanetti MS, Zoller S, Anisimova M. 2013. CodonPhyML: Fast Maximum Likelihood Phylogeny Estimation under Codon Substitution Models. Mol. Biol. Evol.:mst034.

Goldman N, Yang Z. 1994. A codon-based model of nucleotide substitution for protein-coding DNA sequences. Mol. Biol. Evol. 11:725–736.

Haynes BF, Kelsoe G, Harrison SC, Kepler TB. 2012. B-cell-lineage immunogen design in vaccine development with HIV-1 as a case study. Nat. Biotechnol. 30:423–433.

Hershberg U, Uduman M, Shlomchik MJ, Kleinstein SH. 2008. Improved methods for detecting selection by mutation analysis of Ig V region sequences. Int. Immunol. 20:683–694.

Hoehn KB, Fowler A, Lunter G, Pybus OG. 2016. The Diversity and Molecular Evolution of B-Cell Receptors during Infection. Mol. Biol. Evol.:msw015.

Huelsenbeck JP, Rannala B. 1997. Phylogenetic Methods Come of Age: Testing Hypotheses in an Evolutionary Context. Science 276:227–232.

Kepler TB. 2013. Reconstructing a B-cell clonal lineage. I. Statistical inference of unobserved ancestors. F1000Research [Internet] 2. Available from: http://www.ncbi.nlm.nih.gov/pmc/articles/PMC3901458/

Kepler TB, Munshaw S, Wiehe K, Zhang R, Yu J-S, Woods CW, Denny TN, Tomaras GD, Alam SM, Moody MA, et al. 2014. Reconstructing a B-cell clonal lineage. II. Mutation, selection, and affinity maturation. B Cell Biol. 5:170.

Lefranc M-P, Lefranc G. 2001. The Immunoglobulin FactsBook. Academic Press

Liao H-X, Lynch R, Zhou T, Gao F, Alam SM, Boyd SD, Fire AZ, Roskin KM, Schramm CA, Zhang Z, et al. 2013. Co-evolution of a broadly neutralizing HIV-1 antibody and founder virus. Nature 496:469–476.

McCoy CO, Bedford T, Minin VN, Bradley P, Robins H, Matsen FA. 2015. Quantifying evolutionary constraints on B-cell affinity maturation. Philos. Trans. R. Soc. B Biol. Sci. 370:20140244.

Peled JU, Kuang FL, Iglesias-Ussel MD, Roa S, Kalis SL, Goodman MF, Scharff MD. 2008. The Biochemistry of Somatic Hypermutation. Annu. Rev. Immunol. 26:481–511.

Rogozin IB, Diaz M. 2004. Cutting Edge: DGYW/WRCH Is a Better Predictor of Mutability at G:C Bases in Ig Hypermutation Than the Widely Accepted RGYW/WRCY Motif and Probably Reflects a Two-Step Activation-Induced Cytidine Deaminase-Triggered Process. J. Immunol. 172:3382–3384.

Smith DS, Creadon G, Jena PK, Portanova JP, Kotzin BL, Wysocki LJ. 1996. Di-and trinucleotide target preferences of somatic mutagenesis in normal and autoreactive B cells. J. Immunol. 156:2642–2652.

Sok D, Laserson U, Laserson J, Liu Y, Vigneault F, Julien J-P, Briney B, Ramos A, Saye KF, Le K, et al. 2013. The Effects of Somatic Hypermutation on Neutralization and Binding in the PGT121 Family of Broadly Neutralizing HIV Antibodies. PLoS Pathogs. [Internet] 9. Available from: http://www.ncbi.nlm.nih.gov/pmc/articles/PMC3836729/

Wei L, Chahwan R, Wang S, Wang X, Pham PT, Goodman MF, Bergman A, Scharff MD, MacCarthy T. 2015. Overlapping hotspots in CDRs are critical sites for V region diversification. Proc. Natl. Acad. Sci. 112:E728–E737.

Wu X, Zhang Z, Schramm CA, Joyce MG, Do Kwon Y, Zhou T, Sheng Z, Zhang B, O’Dell S, McKee K, et al. 2015. Maturation and Diversity of the VRC01-Antibody Lineage over 15 Years of Chronic HIV-1 Infection. Cell 161:470485.

Yaari G, Benichou JIC, Vander Heiden JA, Kleinstein SH, Louzoun Y. 2015. The mutation patterns in B-cell immunoglobulin receptors reflect the influence of selection acting at multiple time-scales. Philos. Trans. R. Soc. B Biol. Sci. 370:20140242.

Yaari G, Uduman M, Kleinstein SH. 2012. Quantifying selection in high-throughput Immunoglobulin sequencing data sets. Nucleic Acids Res.:gks457.

Yaari G, Vander Heiden JA, Uduman M, Gadala-Maria D, Gupta N, Stern JNH, O’Connor KC, Hafler DA, Laserson U, Vigneault F, et al. 2013. Models of Somatic Hypermutation Targeting and Substitution Based on Synonymous Mutations from High-Throughput Immunoglobulin Sequencing Data. Front. Immunol. [Internet] 4. Available from: http://www.ncbi.nlm.nih.gov/pmc/articles/PMC3828525/

Yang Z. 2007. PAML 4: Phylogenetic Analysis by Maximum Likelihood. Mol. Biol. Evol. 24:1586–1591.

Ye J, Ma N, Madden TL, Ostell JM. 2013. IgBLAST: an immunoglobulin variable domain sequence analysis tool. Nucleic Acids Res. 41:W34–W40.

Zhang W, Bardwell PD, Woo CJ, Poltoratsky V, Scharff MD, Martin A. 2001. Clonal instability of V region hypermutation in the Ramos Burkitt’s lymphoma cell line. Int. Immunol. 13:1175–1184.

